# A Minimally Perturbative DARPin Probe for Quantitative Fluorescence Imaging of the Human TCR-CD3 Complex

**DOI:** 10.64898/2026.09.24.754129

**Authors:** Martí Pons Romero, Barbora Kalousková, Jovana Kloimwieder, Alejandro Linares, Anežka Májková, Lukas Riedl, Caroline Kopittke, Marina Bishara, Mario Brameshuber, Eva Sevcsik

## Abstract

Fluorescence microscopy is a powerful tool for dissecting the molecular mechanisms of T-cell antigen recognition in living cells, but its quantitative insight critically depends on non-perturbative, high-quality probes. Here, we repurpose a small (∼15 kDa) CD3ε-binding DARPin (designed ankyrin repeat proteins) to a fluorescent label for T-cell receptor (TCR)/CD3 complexes on primary human CD8^+^ T-cells, with the aim of generating a powerful tool for quantitative analysis, single-molecule tracking, and advanced imaging of TCR dynamics. We show that the DARPin binds CD3ε with high affinity and selectivity and using single molecule tracking and brightness analysis, we characterize the TCR-CD3 diffusion behavior and show that the DARPin binds to both CD3ε subunits. Importantly, labeling preserves antigen sensitivity: on supported lipid bilayers presenting cognate pMHC, T-cells remain responsive, assemble synapses, form TCR microclusters, and initiate signaling similar to unlabeled controls. We further demonstrate compatibility with lattice light-sheet microscopy for volumetric imaging of T-cell–APC interactions in living cells. Together, these results establish DARPins as versatile, minimally perturbative probes for high-resolution, quantitative studies of T-cell synapse organization and signaling.

## INTRODUCTION

T-cells discriminate foreign from self with extraordinary sensitivity and selectivity, detecting a few or even a single agonist peptide-MHC (pMHC) complexes among thousands of self-ligands on the surface of antigen-presenting cells (APCs) [1,2]. This performance is mediated by the T-cell receptor (TCR)/CD3 complex, whose engagement of pMHC initiates the adaptive cellular immune response. How this precise recognition is achieved at the molecular level remains incompletely understood but likely emerges from the concerted interplay of receptor-ligand binding kinetics, mechanical forces, intracellular signal amplification, and the spatial organization of the immune synapse organization [3,4].

Fluorescence microscopy - often in combination with supported lipid bilayers (SLBs) that reconstitute key features of the APC membrane - has been instrumental in dissecting the molecular mechanisms of T-cell antigen recognition[5]. Using labeled TCR-CD3 and associated reporters, early live-cell imaging established that upon pMHC engagement, TCRs coalesce into microclusters, recruit proximal signaling molecules, and undergo actin-driven centripetal transport to assemble - with adhesion and co-stimulatory receptors - the supramolecular activation clusters (SMACs) of the immune synapse [6–8]. Subsequent single-molecule approaches resolved TCR-pMHC interaction dynamics via *in situ* Förster resonance energy transfer (FRET) measurements reporting TCR-pMHC binding lifetimes[9], or long-illumination single-molecule tracking of fluorescent pMHC on SLBs [10]. Super-resolution imaging mapped the nanoscale organization of the TCR and associated signaling molecules [11–14], and lattice light-sheet microscopy visualized immune synapse remodeling in three dimensions over extended timescales [15,16]. Collectively, these studies have revealed that T-cell activation is governed not only by the biochemistry of individual receptor-ligand interactions, but also by the spatial and temporal coordination of signaling at the nanometer to micrometer scale - underscoring the central role of advanced fluorescence microscopy in uncovering the mechanistic logic of antigen discrimination.

The performance of such fluorescence-based approaches critically depends on the molecular probes used, with requirements varying by application. For fixed-cell imaging, conventional antibodies are often sufficient, but live-cell studies demand probes that minimally perturb the system under investigation. Monovalent Fab fragments against TCR-CD3 have been employed to avoid receptor cross-linking and activation artefacts [10,17]. However, Fab fragments derived from anti-CD3ε antibodies such as OKT3 and UCHT1 can stabilize an active CD3ε conformation and promote Nck recruitment even in the absence of cross-linking [18], potentially perturbing the signaling processes under investigation. Their relatively large size (∼50 kDa) further limits epitope accessibility and spatial precision. A smaller alternative, the H57-derived single chain antibody fragment (scF_V_) (∼27 kDa) against the murine TCR, has enabled single-molecule FRET measurements of TCR-pMHC binding dynamics in living T-cells [9], but no equivalent probe exists for the human TCR. Nanobodies would offer a comparable size advantage and have been used effectively in super-resolution microscopy of other targets [19], but analogous probes for non-perturbative imaging of the human TCR-CD3 complex are lacking.

Here, we address this gap by implementing a DARPin (designed ankyrin repeat proteins)-based probe (∼15 kDa) for quantitative fluorescence imaging of the TCR in primary human CD8⁺ T-cells. DARPins are small (∼14–18 kDa), monomeric, cysteine-free proteins built from stacked ankyrin repeat modules that form a rigid, concave binding surface[20,21]. They can be selected from large combinatorial libraries by ribosome display to bind virtually any target with affinities rivalling those of antibodies. Their high expression yields in *E. coli*, thermal stability, and tolerance of site-specific chemical conjugation make them well suited for fluorescence imaging, while their small size minimizes both linkage error in super-resolution microscopy and steric perturbation in crowded molecular environments [22]. Despite these favorable properties and their widespread use as therapeutic scaffolds and *in vivo* imaging agents [21,23] DARPins have rarely been employed in fluorescence microscopy - limited to a single report of intracellular actin labeling [24] - and have not, to our knowledge, been applied as quantitative imaging probes for cell surface receptors.

A CD3ε-binding DARPin domain has recently been developed and refined as part of bi- and trispecific T-cell engagers and binds CD3ε with high affinity (K_D_ = 6 nM by SPR) [25,26]. Here, we repurpose this domain as a fluorescent probe (∼15 kDa) for quantitative imaging of the TCR-CD3 complex on primary human CD8⁺ T-cells expressing the RA14 T-cell receptor. We systematically evaluate its binding properties, receptor occupancy, and the impact of DARPin labeling on T-cell activation. Combining flow cytometry, single-molecule microscopy, and 3D live-cell imaging, we demonstrate the applicability of the probe across multiple fluorescence imaging modalities. Using SLBs presenting pMHC and ICAM-1 as a minimal model of the APC interface, we show that DARPin-labeled T-cells retain their exceptional antigen sensitivity while enabling quantitative analysis of TCR diffusion, microcluster formation, and immune synapse dynamics. Together, these results establish DARPins as versatile, minimally perturbative probes for quantitative studies of TCR organization and antigen recognition.

## RESULTS AND DISCUSSION

### DARPin expression and TCR binding characterization

To generate a minimally perturbative probe for TCR labeling on human T-cells, we employed a DARPin targeting the CD3ε subunit of the TCR-CD3 complex based on an amino acid sequence reported by Venetz-Arenas et al. [25]. The DARPin (∼15 kDa) was expressed in *E. coli* BL21 (DE3) from a pET vector encoding the DARPin sequence extended with a C-terminal cysteine for maleimide-based fluorophore conjugation and a StrepTag for purification (see Methods, Supplementary Figure 1*a*). Structural modeling using AlphaFold2 Multimer [27,28] predicted a compact architecture comprising three ankyrin repeat domains with a concave binding surface engaging the CD3ε ectodomain (Figure 1*a*, panel i). For fluorescence applications, the DARPin was site-specifically conjugated to AF647 (Lumiprobe, DARPin-AF647) or Cy3B (Lumiprobe, DARPin-Cy3B) with a degree of labeling (DOL) of 0.8 and 0.5, respectively. For reference, Figure. 1*a*, panel ii shows a model of the full TCR-CD3 complex (PDB: 9IRS) with DARPins placed on both CD3ε subunits, and Figure 1*a*, panel iii a CD3δε heterodimer complexed with an OKT3 Fab fragment, illustrating the relative probe dimensions.

**Figure 1.**
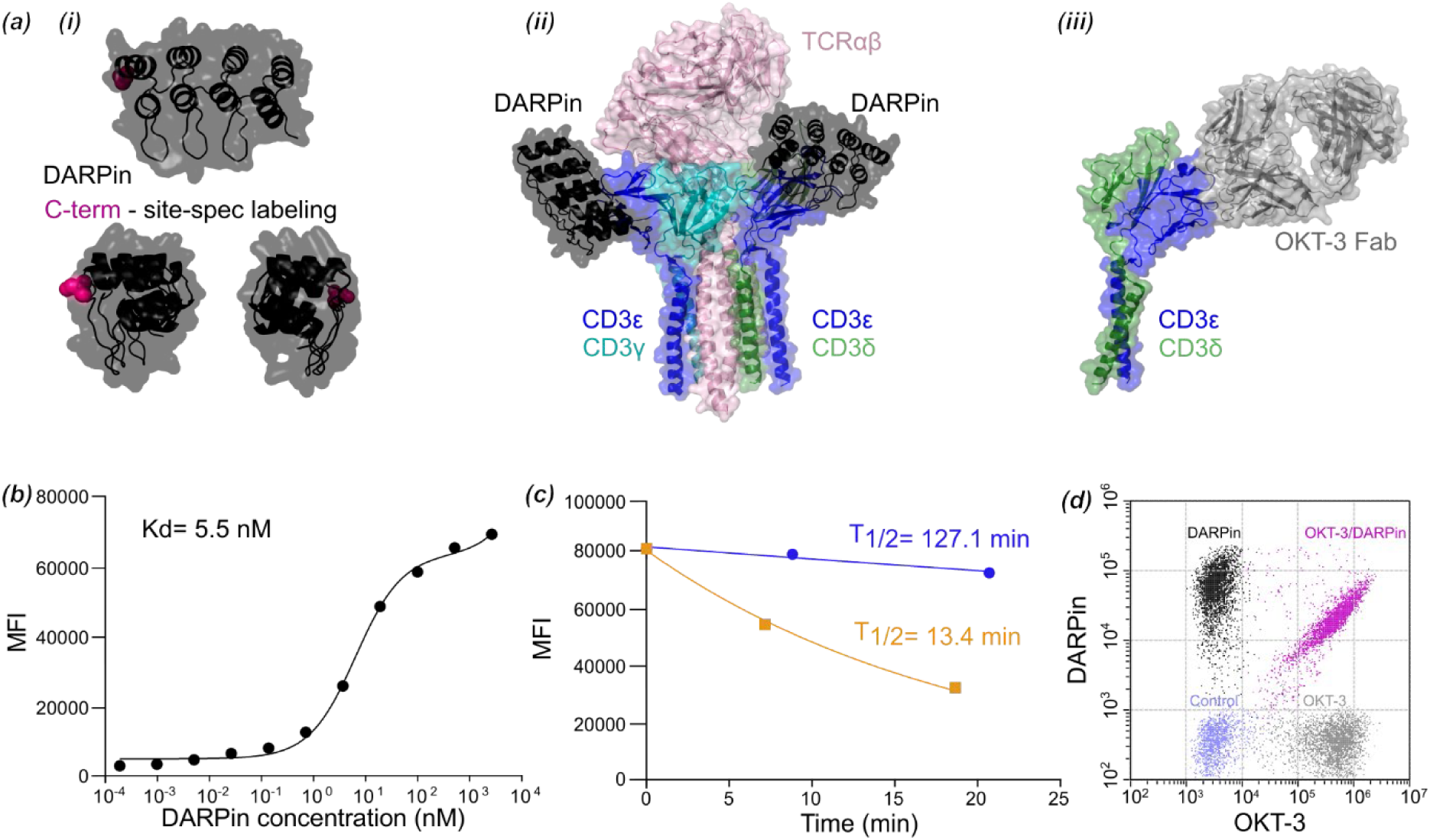
Development and biochemical characterization of a fluorescently labeled TCR-CD3-binding DARPin. (*a*) Structural model of the TCR-CD3-binding DARPin and site-specific fluorophore labeling strategy (i). CD3ε subunits with bound DARPin in context of the whole TCR-CD3 complex (ii) and CD3δε with OKT3 Fab (iii). (*b*) Binding of the DARPin to T-cells expressing the human TCR-CD3 complex, determined by flow cytometry. (*c*) Dissociation kinetics of DARPin binding to the TCR-CD3 complex at 4°C (blue) and room temperature (orange). (*d*) Competition of DARPin binding with the anti-CD3 antibody OKT3. For reference, T-cells were stained with either OKT3 antibody (grey population) or the DARPin (dark population). The binding of DARPin after pre-incubation with OKT3 is reduced due to competition for the same epitope (pink population). However, due to its small size, the DARPin can still bind to non-occupied epitopes. Control cells were incubated with a non-binding secondary antibody (blue population).

To characterize DARPin binding to TCR-CD3 complexes, we performed a DARPin-AF647 titration on RA14 CD8⁺ T-cells by flow cytometry. The DARPin showed robust, concentration-dependent staining with a binding curve that plateaued at approximately 200 nM (Figure 1*b*), indicating specific, saturable binding to surface-expressed CD3ε. This was used as the standard labeling concentration for all subsequent experiments. Negligible binding was observed on CD3ε-negative control cells, confirming target specificity (Supplementary Figure 1*b*). To map the DARPin binding site relative to established anti-CD3ε antibodies, we performed competition experiments with OKT3. Pre-incubation with OKT3 in saturation (2 µg/ml) reduced subsequent DARPin binding substantially, consistent with an overlapping or proximal epitope on CD3ε [26] (Figure 1*d*). Notably, the DARPin labeled a larger fraction of surface CD3ε than the full-size OKT3 antibody (∼150 kDa), suggesting that its smaller size (∼15 kDa) allows it to access CD3ε sites within the TCR–CD3 complex that are sterically occluded for the antibody.

### DARPin-based labeling enables quantitative TCR imaging

We next assessed DARPin-based labeling in fluorescence microscopy experiments. RA14 CD8⁺ T-cells stained with DARPin were imaged on SLBs under non-activating conditions (ICAM-1 only) using total internal reflection fluorescence (TIRF) microscopy (Figure 2*a*). Quantification of the DARPin surface density yielded an average of ∼71 ± 6.2 molecules/µm² (mean ± SEM) for RA14 T-cells (Figure 2*b*). On the same non-activating surfaces, we characterized the diffusion behavior of DARPin-labeled TCR– CD3 complexes by single-molecule tracking. TCR–CD3 exhibited Brownian-like motion with a mean diffusion coefficient of D = 0.1 ± 0.004 µm²/s (Figure 2*c*). These values are in close agreement with the diffusion coefficient of D = 0.082 µm²/s reported for HaloTag-labeled 1G4 TCR in JurkaT-cells [17], indicating that DARPin binding does not significantly alter TCR lateral mobility. Notably, in the same study, TCRs labeled with a UCHT1 Fab fragment diffused substantially more slowly (D = 0.028 µm²/s) - approximately threefold reduced compared to the unperturbed receptor. While the larger size of the Fab (∼50 kDa vs. ∼15 kDa for the DARPin) may contribute through increased hydrodynamic drag, the dominant effect is likely functional: UCHT1 Fab binding induces a conformational change in CD3ε that exposes its proline-rich sequence and recruits the cytoskeletal adaptor Nck [18], coupling the receptor to the actin cytoskeleton and thereby reducing its lateral mobility. That the DARPin preserves high TCR diffusion suggests it does not trigger this conformational switch.

**Figure 2.**
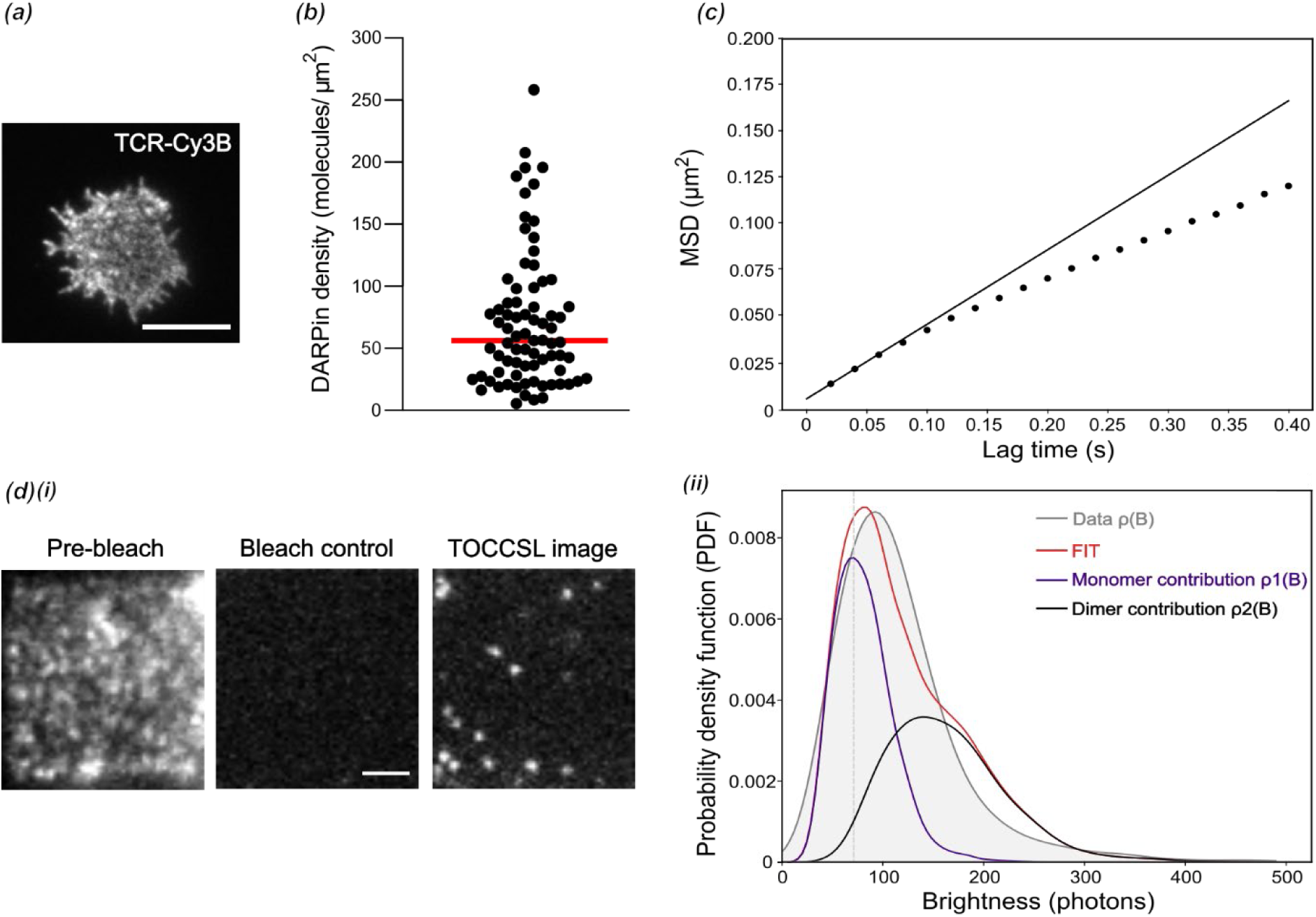
DARPin-based labeling enables quantitative analysis of TCR density, mobility, and stoichiometry in primary human T-cells. Representative TIRF image of a DARPin-Cy3B-labeled RA14 T-cell, recorded 5 min after initial contact. Scale bar, 10 µm. (*b*) Surface density of DARPins on RA14 T-cells. The median of 56 molecules/µm^2^ is indicated in red. (*c*) Mean square displacement (MSD) of DARPin-AF647-labeled TCRs as a function of lag time. The diffusion coefficient was determined by fitting Eq. 6 (black line). Only the first two data points were considered for the fit. (d) TOCCSL approach. (*i*) A defined region of the basal T-cell membrane was illuminated in TIR mode through an aperture in the excitation pathway (pre-bleach image). A high-intensity bleach pulse abolished all fluorescent signals within the region of interest (bleach control image). After a recovery interval of 15 s, DARPin-labeled TCR–CD3 complexes that had diffused in from the surrounding unbleached membrane were recorded as isolated diffraction-limited spots (TOCCSL image). Scale bar, 2 µm. (*ii*) Brightness distributions ρ_n_ of localizations detected in the TOCCSL image are plotted, fitted and deconvolved into monomer and dimer contributions to reveal 49% ± 1 % dimers for DARPin-AF647 (n = 79 cells).

The TCR-CD3 complex contains two CD3ε chains in distinct molecular contexts, one in the CD3γε heterodimer and the other in the CD3δε heterodimer, thus the DARPins could, in principle, bind both sites. For a quantitative evaluation of the labeling stoichiometry of the TCR-CD3 complex, we resorted to brightness analysis of individual TCR-CD3 complexes. Because of the high surface density of TCR-CD3 on resting T-cells, individual receptors cannot be resolved by diffraction-limited microscopy. We therefore used TOCCSL (Thinning Out Clusters While Conserving Stoichiometry of Labeling [29], Figure 2*d*, panel i) analysis on living T-cells labelled with DARPin-AF647. For this, a defined region of the basal membrane was photobleached by a short, high-intensity laser pulse, and after a recovery interval (15 s), as unbleached DARPin-AF647-labeled TCR–CD3 complexes diffused into the bleached area, isolated fluorescent signals were recorded. We compared the measured distribution of fluorescence brightness values ρ(B) with the brightness distribution of single DARPin-AF647, ρ_1_, which was obtained recording T-cells with 80-fold substoichiometric TCR-CD3 labeling (Methods section). A weighted linear combination of n-mer contributions ρₙ derived from ρ₁ was fitted to the measured brightness distribution ρ(B) (Methods, Eq. 5), allowing classification into populations carrying one or two DARPins per TCR–CD3 complex (Figure 2*d*, panel ii). Importantly, when T-cell populations were stratified by low, medium, and high DARPin labeling density and analyzed separately, the dimer fraction remained consistent across all groups (Supplementary Figure 2). This indicates that cells with lower DARPin staining carry fewer TCR–CD3 complexes on their surface, rather than having experienced probe dissociation during the experiment. Assuming binomial labeling statistics, the observed dimer fraction of ∼50% corresponds to ∼70% of TCR–CD3 complexes carrying two DARPins after correcting for a labeling efficiency of 0.8., indicating that both CD3ε subunits are accessible. This likely represents an underestimation of the true fraction of doubly accessible complexes, as incomplete photobleaching due to diffusion of complexes during the photobleaching pulse and an imperfect laser bleach profile reduces the probability of detecting two fluorophores even when both binding sites are occupied [30]. Taking this into account, the average TCR surface density on RA14 T-cells can be estimated to be between 44 and 52 TCRs/µm^2^.

### DARPin labeling preserves antigen-dependent TCR reorganization and signaling

To verify that DARPin labeling neither activates T cells nor alters their antigen sensitivity, we performed a series of functional assays. We compared TCR dynamics on SLBs functionalized with Alexa Fluor 647–labeled pMHC (HLA-A*02:01/CMV) and ICAM-1 (activating conditions) versus ICAM-1 alone (non-activating conditions). Under non-activating conditions, cells spread actively on the bilayer, extending highly motile filopodia, while TCRs were randomly distributed (Figure 3*a*, Supplementary movie 1). At a density of ∼ 10 pMHC /µm², TCRs formed microclusters that coalesced into a central supramolecular activation cluster (cSMAC) within 5-10 minutes, consistent with immune synapse formation (Figure 3*a*, Supplementary movie 2 and 3). These findings confirm that TCR microcluster and cSMAC formation is strictly antigen-dependent and that the DARPin label does not induce artifactual clustering.

**Figure 3.**
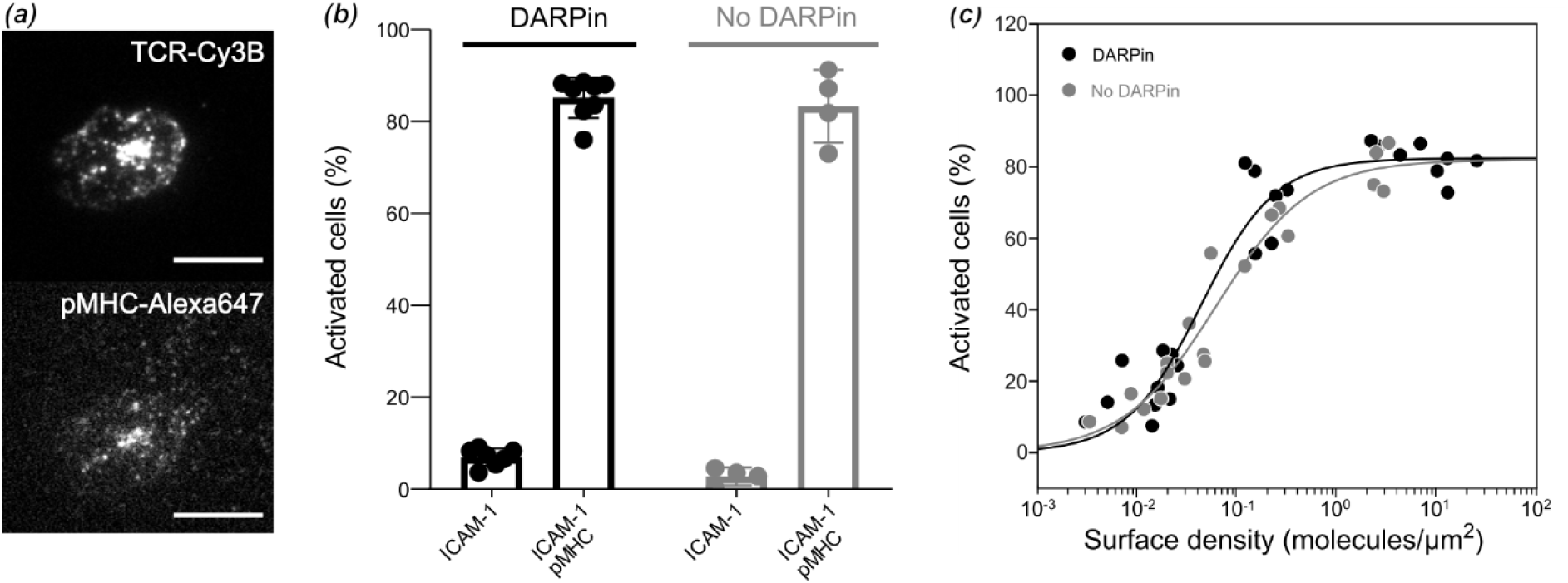
DARPin labeling preserves T-cell activation and effector function. (*a*) Representative TIRF image showing the formation of DARPin-Cy3B-labeled TCR microclusters and their co-localization with Alexa647-pMHC at a density of ∼10 molecules/µm². The image was recorded 5 min after first contact. Scale bar, 10 µm. (*b*) Percentage of T-cells showing calcium influx. The influence of DARPin labeling at saturating conditions is compared to control cells under non-activating (ICAM-1 only) and activating conditions (ICAM-1 + pMHC). (*c*) Dose–response curves for T-cell activation at different pMHC surface densities. Each data point corresponds to the percentage of activated cells determined in an individual experiment at a specific pMHC density. Fitting of dose– response curves with Eq. 3 (see Methods) yielded the activation thresholds in the absence (0.06 ± 0.02 pMHC/ µm^2^) and presence (0.04 ± 0.01 pMHC/µm^2^) of DARPin labeling. Activation curves of DARPin-labeled (black) and unlabeled cells (grey) are compared.

Next, we evaluated whether DARPin labeling interferes with T-cell activation using ratiometric calcium imaging. Labeling neither triggered activation under non-activating conditions (ICAM-1 only) nor impaired activation under activating conditions (ICAM-1 + pMHC) (Figure 3*b*). To assess whether DARPin-AF647 labeling alters antigen sensitivity, dose–response curves were recorded across different pMHC surface densities on SLBs in the absence or presence of DARPin labeling. Activation thresholds and maximal responses were similar between conditions, with no significant differences in EC_50_ (Figure 3*c*), indicating that DARPin labeling fully preserves T-cell activation.

To demonstrate compatibility of DARPin labeling with volumetric live-cell imaging, we performed lattice light-sheet microscopy (LLSM) of DARPin-labeled RA14 CD8⁺ T-cells interacting with K562 feeder cells (Figure 4). This imaging modality provides diffraction-limited 3D resolution across entire cell volumes with minimal phototoxicity, enabling extended observation of dynamic cell-cell interactions. T-cells labeled with DARPin-AF647 were pipetted together with unlabeled antigen-presenting K562 cells and imaged at room temperature, acquiring sample scans every 60 or 120 seconds for total durations of up to 10 minutes. Time-lapse LLSM of living cells captured the spatiotemporal dynamics of T-cell–feeder cell engagement under both non-stimulatory and activating conditions. In the absence of cognate antigen, T-cells actively scanned the feeder cell surface (Figure *4a*). Under activating conditions, T-cells polarized toward the feeder cell, formed stable contacts, and accumulated TCR at the cell-cell interface into a cSMAC-like structure (Figure 4*b*). The DARPin label produced bright, specific TCR signal throughout the T-cell surface with no detectable background on the feeder cells, confirming that DARPin labeling is fully compatible with 3D imaging of TCR organization during physiological T-cell-APC interactions.

**Figure 4.**
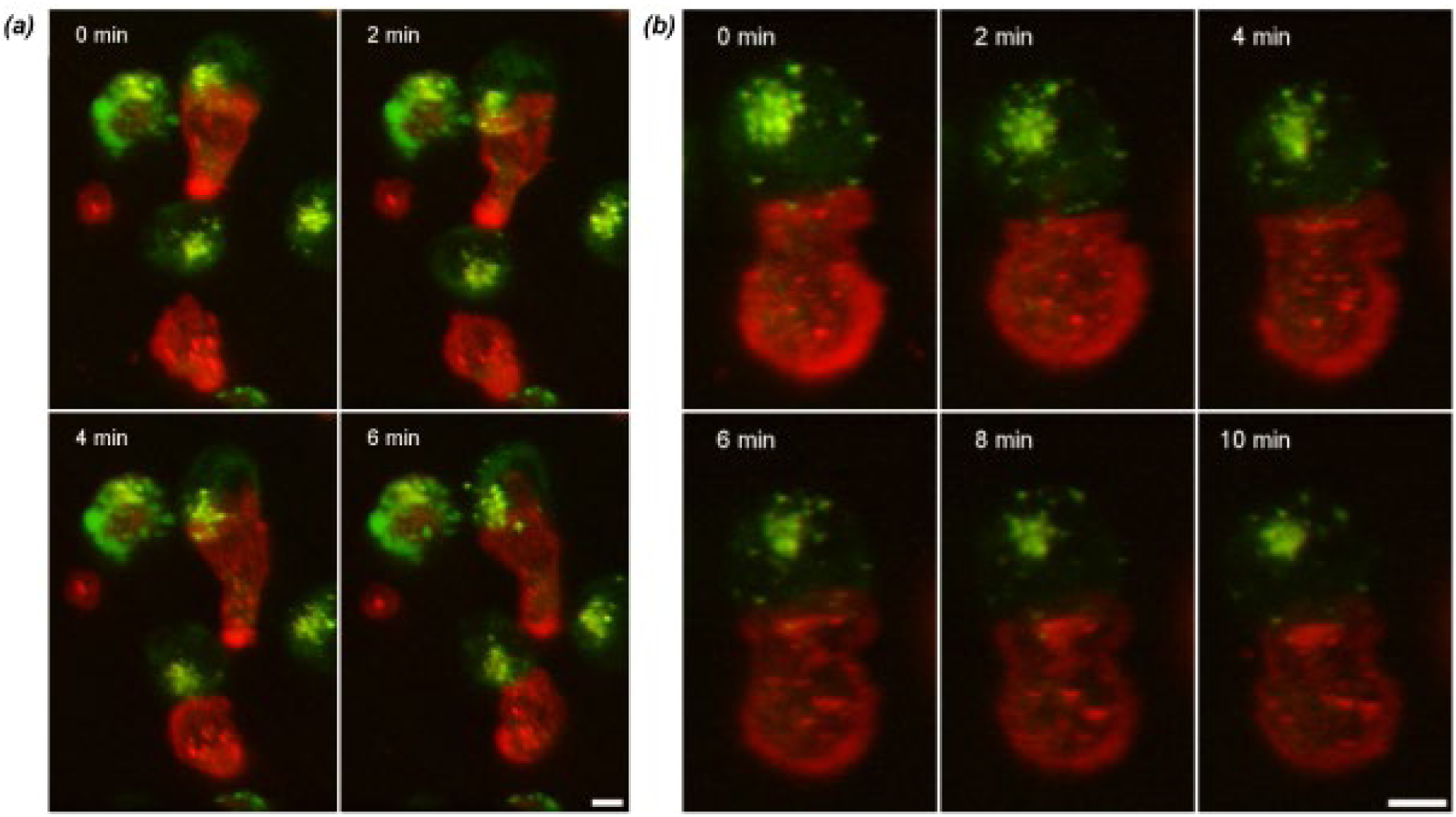
DARPin labeling is compatible with lattice light-sheet imaging of T-cells. LLSM images of DARPin-AF647-labeled RA14 T-cells (TCR–DARPin, red) interacting with K562 feeder cells (autofluorescence, green) in the absence (*a*) and presence (*b*) of cognate peptide. Shown are select time points of deskewed volumetric sample scans taken every 120 s (*a*) or every 60 s (*b*) in 3D maximum rendering. Both time series were acquired at room temperature. Scale bar, 5 µm.

## CONCLUSION

We introduce a compact, DARPin-based fluorescent probe (∼15 kDa) that addresses a practical gap in the current toolkit for imaging the human TCR–CD3 complex. The probe engages CD3ε specifically within the TCR–CD3 complex, and labels both CD3ε subunits. Labeling is consistent with non-perturbative behavior: receptor mobility remains rapid and antigen sensitivity is not detectably altered. The probe also supports robust volumetric 3D visualization of TCR organization during T-cell-APC encounters. Together, these results support the DARPin as a minimally perturbative label suitable for quantitative measurements of receptor stoichiometry, dynamics, and signaling, with practical advantages of small size, monovalency, and site-specific conjugation.

The DARPin platform is readily extensible. The same expression and site-specific conjugation strategy can be adapted to other (immune) receptors, enabling multicolor, multiplexed measurements of receptor crosstalk at minimal steric cost. The probe’s size and rigidity should be advantageous for high-precision applications such as DNA-PAINT, MINFLUX, expansion microscopy, and correlative fluorescence-electron microscopy. More broadly, our findings highlight DARPins as an underused class of quantitative imaging probes for cell biology, bridging the gap between nanobody-scale labels and antibody-derived reagents for non-perturbative interrogation of receptor organization and antigen recognition.

## MATERIALS AND METHODS

### Structural model

The structure of the DARPin alone or in complex with the CD3εγ (sequence taken from PDB:1SY6 [31], chain C) and CD3εδ (sequence from PDB:1XIW [32], chains A and B) heterodimer was predicted using AlphaFold-Multimer v3 [28] via ColabFold [33]. Structures were visualized in PyMOL (PyMOL Molecular Graphics System, Version 3.1, Schrödinger, LLC).

### DARPin protein expression

Nucleotides encoding the DARPin amino acid sequence (version 3 of the CD3ε-specific DARPin [25]; MDLGQKLLEAAWAGQLDEVRILLKAGADVNAKNSRGWTPLHTAAQTGHLEIFEVLLKAGA DVNAKTNKRVTPLHLAAALGHLEIVEVLLKAGADVNARDTWGTTPADLAAKYGHRDIAEV LQKAA), extended with a short flexible linker, a cysteine for site-specific labeling and a Strep-tag (extension sequence GSCGSWSHPQFEK), were cloned into the pET-30a(+) expression vector. *E. coli* BL21(DE3) transformed with this plasmid were cultured overnight (50 ml LB medium, 50 µg/ml kanamycin, 37 °C, 230 rpm). This starter culture was used to inoculate 500 ml LB medium (50 µg/ml kanamycin), which was grown to an OD₆₀₀ of 0.6–0.8 at 30 °C and 230 rpm. Protein expression was then induced with IPTG (0.5 mM final concentration) and continued for 16 h under the same conditions. Cells were harvested by centrifugation the following day, and the pellet was stored at −20 °C.

The protein was purified using Strep-Tactin agarose (Cube Biotech) according to the manufacturer’s protocol. Briefly, the cell pellet corresponding to 200 ml of culture was resuspended in 10 ml lysis buffer (100 mM Tris, 150 mM NaCl, protease inhibitor cocktail (Omega), 1 mg/ml lysozyme, pH 8). The suspension was sonicated on ice and incubated on a shaker for 1 h. The lysate was clarified by centrifugation (15,000 × g, 10 min, 4 °C) and filtered (0.45 µm). Strep-Tactin agarose was equilibrated in wash buffer (100 mM Tris, 150 mM NaCl, pH 8), and 0.5 ml of the slurry was added to the lysate and incubated for 2 h. The resin was then transferred to a gravity-flow column and washed three times with 5 ml wash buffer. The protein was eluted with elution buffer (100 mM Tris, 150 mM NaCl, 2.5 mM desthiobiotin, pH 8) and dialyzed overnight against 0.5 l PBS (pH 7.4) using a 3 kDa MWCO dialysis cassette. Protein quality was verified by SDS-PAGE.

For labeling, the protein was first incubated with 5 mM TCEP for 2 h at 4 °C, after which a 20-fold molar excess of maleimide-conjugated fluorophore (Cy3B or AF647; both Lumiprobe) was added, and the reaction was continued overnight at 4 °C. Unreacted dye was removed using an Amicon concentrator. Protein concentration and degree of labeling (DOL) were determined using a NanoDrop spectrophotometer.

### Cell culture

Antigen-specific CD8⁺ T-cells expressing the RA14 TCR, which recognizes the CMV pp65 peptide presented by HLA-A2/CD80-engineered K562 target-cells, were generated essentially as described previously [34,35]. Briefly, PBMCs were isolated from human buffy coats (healthy donors, Red Cross) and sorted for CD8⁺ cells. These cells were expanded using a T-cell-specific activator (ImmunoCult Human CD3/CD28 T Cell Activator, Stemcell) and IL-2 (Miltenyi Biotec). Blasting cells were then electroporated to knock out the endogenous TCR and to introduce the RA14 TCR sequence using a CRISPR/Cas9 strategy. After recovery, cells were labeled with HLA-A2/CMV tetramers to identify RA14-positive cells, which were sorted by FACS and subsequently expanded in an antigen-dependent manner using feeder cells (HLA -A2/CD80-engineered K562 targets pulsed with CMV pp65 peptide) [36]. After two rounds of expansion, the cells were frozen. Prior to experiments, T-cells were thawed and challenged once more with antigen-presenting feeder cells. Experiments were performed 7–10 days after activation. The T-cell co-culture was maintained in ImmunoCult-XF T-cell Expansion Medium (Stemcell) supplemented with 50 U/mL IL-2 and Pen/Strep. Feeder cells were kept in PRMI medium supplemented with 10% FBS, Pen/Strep.

### Flow cytometry

Binding curves, half-lives, and competition with the OKT3 monoclonal antibody were assessed by flow cytometry on a CytoFLEX SRT (Beckman Coulter), using CytExpert software (version version 1.2.0.10004) for both acquisition and analysis. To determine the binding properties of the DARPin, RA14 T-cells were harvested and aliquoted at 200,000 cells per condition. Cells were incubated with fluorescently labeled DARPin at the indicated concentrations for 30 min on ice, washed twice with HBSS, and kept on ice until analysis. For half-life determination, cells were labeled with DARPin at a saturating concentration (4 µg/mL) as above and then half of the sample was maintained at room temperature; the decay of bound fluorescence was monitored over time. To assess competition with OKT3, cells were first incubated with unlabeled OKT3 antibody (2 µg/mL; BioLegend, cat. 317302, lot B231404), washed, and then incubated with labeled DARPin (4 µg/mL) together with a secondary antibody (goat anti-mouse IgG, Abberior STAR 488, 10 µg/mL; cat. ST488-1001-500UG, lot 01201PK-1). Mean fluorescence intensities (MFI) were plotted in GraphPad Prism (version 10.6.1). Equilibrium dissociation constants (K_d_) were obtained by fitting a one-site total binding binding model,

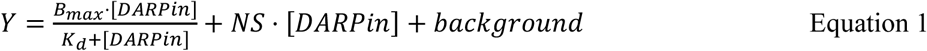

where Y is the MFI, B_max_ the maximal binding, and K_d_ the DARPin concentration at half-maximal binding, NS is the slope of nonspecific binding in Y units divided by X units, background is the baseline.

Half-lives (*t₁/₂*) were obtained by fitting a one-phase exponential decay model,

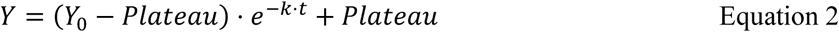

where *Y_0_*is the MFI at *t* = 0, Plateau the MFI at late time points (here value of unlabeled cells), *k* the decay rate constant, and *t* time.

### Vesicles and bilayer preparation

SLBs were essentially made as described [37]. Briefly, vesicles containing 98% POPC (1-palmitoyl-2-oleoyl-sn-glycero-3-phosphocholine; Avanti Polar Lipids, 850475P) and 2% DGS-NTA(Ni) (1,2-dioleoyl-sn-glycero-3-[(N-(5-amino-1-carboxypentyl)iminodiaceticacid) succinyl] nickel salt; Avanti Polar Lipids, 790404P) were prepared as described [38]. Glass slides (#1.5, 25x75 mm, Knittel Glass) were cleaned for 15 min in Piranha cleaning containing concentrated H_2_SO_4_ and 30% H_2_O_2_ (both from Sigma-Aldrich) at a ratio of 3:1 (acid/peroxide) and glued to LabTek 16-well chambers (Thermo Fisher Scientific, 10507401). Vesicles were added immediately and incubated for 15 minutes at room temperature. Bilayers were washed extensively with PBS, which was later exchanged for a solution of 0.1% BSA (bovine serum albumin; Sigma-Aldrich, A3059) in PBS and incubated for 15 minutes. His-tagged proteins were added and incubated for 60 minutes at room temperature.

### Calcium labeling, imaging and analysis

T cells were labelled with 50 µg/ml Fura-2 AM (Invitrogen, F1221) for 15 minutes at room temperature and then washed with HBSS supplemented with 2% fetal bovine serum (FBS). T cells final concentration was adjusted to 7 × 10^6^cells/ml. Before adding cells, the PBS of bilayers was exchanged for HBSS + 2% FBS. 3 × 10^4^ cells were used per experimental condition at room temperature.

Immediately after, imaging was conducted at an Axiovert 200M microscope (Zeiss), equipped with a Polychrome lightsource (FEI, Till Photonics), a dichroic (Chroma, T400lp), a 10x objective (UplanFLN, NA 0.3; Olympus), an emission filter (Chroma, ET510/80), and a EM-CCD camera (Andor, iXon DU-897). Live Acquisition 2.6.0.12 (FEI) was the software used to control the microscope. Cells were illuminated alternately at 340 nm and 380 nm, with illumination times of 75 ms of 30 ms, respectively, resulting in a frame rate of 1 fps. Ratiometric calcium data was analyzed using Python (3.11.6), with scripts implemented as jupyter (1.0.0) notebooks (7.0.6) and algorithms building on functions from sdt-python [39], a comprehensive analysis package for fluorescence microscopy data. NumPy [40], SciPy [41] and pandas (2.1.2, 10.25080/Majora-92bf1922-00a and 10.5281/zenodo.10045529) supported data treatment and analysis. Trackpy (v0.6.1, [42]) was used for cell tracking. Matplotlib (3.8.1, [43], 10.5281/zenodo.10059757) and seaborn (0.13.0, [44]) facilitated data visualization.

Before analysis, all microscopy records were reviewed and early frames, captured before the final measurement position was found, were excluded. To determine cell positions, we generated sum images for each time point by adding the image pairs from both channels (340nm + 380nm). Cells were localized (sdt.loc.cg.locate(…, radius=3-4, …)), based on [45], using optimal brightness thresholds for each measurement day. For each localization, the background-corrected Fura-2 brightness was determined from the raw images in both channels (sdt.brightness.from_raw_image(…, radius=4, bg_estimator=“mean”, mask=“circle”). Subsequently, Fura-2 ratios were calculated by dividing the cells’ brightness in the 340nm-channel by their brightness in the 380nm-channel.

Cell localizations were filtered for x-y-positions, applying a 341x484 px-ROI that excluded the under-illuminated edges of the microscope’s field of view. The localizations were linked into trajectories (trackpy.link(…, search_range=4, memory=3)) and the momentary speed of each cell was calculated with a rolling window of three frames. Fura-2 ratio and speed curves were smoothed (scipy.signal.savgol_filter(…, window_length=101, polyorder=3). The Fura-2 ratio slope was calculated by deriving the smoothed Fura-2 ratio curve in a window of 1, followed by an additional smoothing step. To prevent overcounting of fractured trajectories, trajectories without a localization in the final frame were discarded. Additionally, trajectory parts where cells still floated above the bilayer (momentary cell speed > 0.213 px /s) were not considered for classification. Finally, trajectories with less than 100 analyzable frames were excluded.

T-cells were classified via two thresholds: (a) the activation threshold (Fura-2 ratio=0.7) to distinguish between cells with high or low intracellular calcium levels and (b) a threshold for the Fura-2 ratio slope to verify that T cells display the steep rise in Fura-2 ratio that is characteristic for calcium influx upon T-cell activation (d/dt Fura-2 ratio = 0.0059/s). T-cell trajectories were sorted into three categories. (1) Non-activating: T-cells with Fura-2 ratios continuously below the activation threshold. (2) No-peak: T-cells with Fura-2 ratios that surpass the activation threshold, but do not show a rising flank, e.g. pre-activated cells. (3) Activating: T-cells that surpass the activation threshold as well as the slope threshold. Percentages of activated T-cells were calculated based on the cell ensemble included in the classification (% of activated cells = number of activated cells / number of (non-activated + activated + no-peak) cells *100).

To obtain a dose-response curve, for each condition, the activated percentage from the calcium classification is complemented with the respective pMHC surface density in molecules/μm^2^. This constructed dose-response curve is fitted using scipy.optimize.curve_fit (Equation 3):

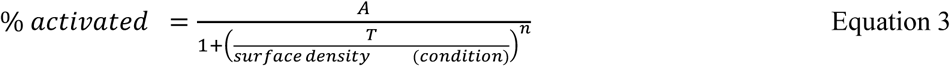

where *A* is the maximal expected response, *T* is the activation threshold (i.e., EC_50_, the surface density for which 50% of cells activate), and *n* is the Hill coefficient. The number of pMHC molecules per μm² on low-density SLBs was determined using the sdt-loc.daostorm 3D algorithm from the sdt_python package [39]. In addition, single pMHC brightness and background were read out with the same algorithm and used to obtain high-occupancy pMHC densities by dividing the background-corrected high-occupancy bulk pMHC brightness by the Alexa647-pMHC single-molecule brightness.

### Total internal reflection fluorescence microscopy (TIRFM)

Fluorescence measurements were essentially performed as described before [37]. TIRFM measurements were acquired on a custom-built microscope based on an inverted Axiovert 200 (ZEISS) equipped with an α Plan-Apochromat 100×/1.46 oil DIC (UV) VIS-IR objective. To excite fluorophores, lasers at wavelengths 532 nm and 642 nm (Oxxius) were used. The excitation beam was deflected away from the optical axis using a mirror mounted on an adjustable piezo-controlled stage to achieve TIR illumination. Excitation and emission were separated using a dichroic mirror (zt532/640/NIR rpc, Chroma F68-532 AHF). An Akrima image splitter was used with two single-band filters (Chroma HQ 585/40M, and Chroma 700/75 ET Bandpass). The emitted signal was projected through a 1× tube lens onto an Andor iXon Ultra EM-CCD camera with 16 μm pixels, operated at -60°C. Timing protocols were developed and performed using an in-house-written package implemented in LABVIEW. Experiments were carried out at room temperature. T cells were recorded simultaneously with the SLBs (pMHC+ICAM-1; Figure 2*a*, Figure 3*a*) using 5 ms of illumination time and a frame rate of 2 fps at laser powers of ∼0.1 kW/cm² (532 nm) and ∼0.2 kW/cm² (642 nm), respectively. For SLBs only (Figure 3*c*) 3ms of illumination time and a frame rate of 100 fps at a laser power of ∼0.8 kW/cm^2^ (642 nm) was used. For TOCCSL (Figure 2d, panel ii), cells were imaged using 5 ms illumination time and a frame rate of 50 fps. The laser power was ∼7.8 kW/cm^2^ (642 nm) for bleaching and ∼0.8 kW/cm² for imaging.

### Single-molecule imaging and analysis (TOCCSL, TCR surface density and mobility)

T-cells were first washed with HBSS and then labeled with 4 µg/ml DARPin (saturating conditions determined via FACS, see Figure 1*b*) or 50 ng/ml (single molecule density) for 30 minutes on ice. Cells were resuspended at a final concentration of 20 × 10^6^cells/ml and kept on ice until use. PBS was exchanged for HBSS for imaging.

8x10^4^ labeled cells were added to the SLB and were left for ∼2 min to settle and spread, followed by imaging for no longer than 15 min. For TOCCSL, a 64x64 pixel (∼100 µm^2^) region of interest (ROI) was defined and centered on the laser’s illumination profile and confined by a rectangular aperture. Measurements consisted of a pre-bleach frame, a bleaching pulse of 1 s, recovery time of 15 s, and subsequent 100 recovery frames at 50 fps with 5 ms exposure time. The recovery time allowed non-bleached molecules from neighboring areas to diffuse into the bleached area and was optimized for high signal yield while preserving well separated signals. Only the first frame was used for brightness analysis.

To determine the brightness of a single fluorophore (monomer reference), T-cells were labeled with DARPin-AF647 at 50 ng/ml. Sequences of 100 images of well-separated mobile single-molecule signals were recorded under the same illumination conditions as TOCCSL recordings, but without bleaching. A custom Python script based on the sdt-python package [39] was used for TOCCSL analysis, which included the detection of fluorescent features, the grouping of cells based on their surface density, filtering, and brightness analysis. The fluorescent features were located using the SDT-python Gui.Locator Plugin by drawing a ROI that included well separated signals in the center of the bleaching region and using the built-in DAOSTORM 3D algorithm (Radius = 1, Model = 2d, Threshold = 100, Max. Interactions = 20, Find-filter = Cg, Min. Distance = 1, Size range = [0.5,2.0]). The localization data was saved, preserving the position, integrated brightness B (mass), full width at half-maximum (FWHM) and local background of fluorescent signals in TOCCSL and monomer reference recordings. Outliers in size and background, which indicate low quality or false positive locations, were excluded from further analysis in the filtering step of the analysis (background: [10,20], size: [0.6,1.4]).

Cells were grouped based on their surface density before bleaching. Surface density was calculated as the ratio of the mean brightness in a centered, rectangular region with half the height and width of the original dimensions of the pre-bleach image and the peak of the probability density function of the single molecule reference. With a grouping threshold of 0.35, low, medium and high expressing cells were separated to find behavioral differences. For the final analysis, all recordings were grouped together (threshold = 1). The brightness analysis to determine the oligomeric composition of TOCCSL data is described in detail in [46]. Signals from the single molecule recordings from images 50-100 in the TOCCSL sequence were used to determine the probability density function (PDF) of monomers, ρ_1_(B). Due to the stochastic photon emission, the PDFs of N colocalized emitters can be calculated by a series of convolution integrals,

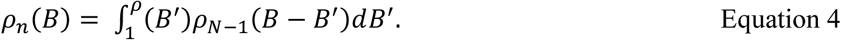

From the known PDFs, a weighted linear combination can then be used to approximate the brightness distribution of a mixed population of monomers, dimers, and higher order oligomers. Brightness values were pooled and used to calculate *ρ*(*B*).

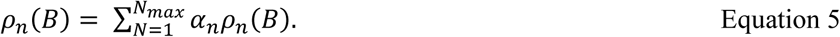

A least-square fit was employed to determine the weights of the individual pdfs, *α_N_*, with 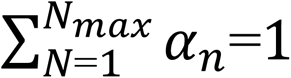 No higher contributions than dimers (*α*_2_) were observed.

To assess TCR surface density and mobility, custom Python scripts based on the sdt_python package [39] were developed and applied to data generated during TOCCSL experiments. First, the single TCR signals were localized from the TOCCSL data (frames 5-105 of each TOCCSL sequence) using the sdt-loc.daostorm 3d algorithm, extracting the total brightness and background of each feature (parameter ‘mass’ and ‘bg‘, respectively), and its x and y coordinates. To define TRC diffusion coefficients, localized single-molecule data were tracked using trackpy [42]. MSD analysis for all trajectories of all cells was performed using sdt.motion.Msd(.., ensemble=True, …), and diffusion coefficient D was calculated with:

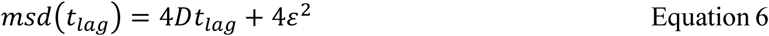

where *ε* is positional accuracy and *t_lall_* is lag time. MSD analysis of individual trajectories sdt.motion.Msd(…, ensemble=False, …). was executed to determine the immobile fraction. To determine TCR surface densities, the mean brightness of the TCR per μm^2^ (*brig*ℎ*tness_BULK_*), obtained from labeling TCR to saturation (frame 1 of each TOCCSL movie), was corrected for the camera offset per μm^2^ (*offset_BULK_*), and divided by the mean single TCR brightness (*brig*ℎ*tness_SM_*), obtained from single TCR localization data:

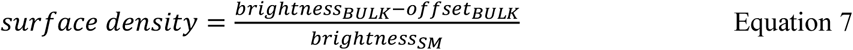

### Lattice Light-Sheet Microscopy

Volumetric imaging was performed on a ZEISS Lattice Lightsheet 7 (LLS7) microscope controlled by ZEISS ZEN software (version 3.13) equipped with dual Hamamatsu ORCA-Fusion sCMOS cameras (6.5 μm pixel pitch; calibrated imaging pixel size 0.145 μm). Excitation used a 13.3×/0.44 NA lattice light-sheet objective (30° relative to the coverslip) and detection used a 44.83×/1.0 NA objective (60° relative to the coverslip). A meniscus lens operated in water-immersion served as the relay optics between sample and detection objective. Built-in free-form optical elements (included in the detection objective) were used to correct for refractive-index mismatches (aberration correction). Samples were mounted in ibidi μ-Slide 8 Well^high^ glass-bottom chambers and placed in the built-in ibidi environmental enclosure on the LLS7 stage; live-cell imaging was performed at room temperature and with all incubation parameters switched off. RA14 CD8^+^ T-cells were purified by Ficoll density gradient centrifugation prior to labeling in saturation with DARPin-AF647 on ice using the previously indicated concentrations and protocol. Freshly labeled and washed cell suspensions (5 Mio./ml) were kept on ice and protected from light before pipetting into the mounted sample chamber (using 2% FBS in HBSS as the imaging buffer) immediately before imaging. As antigen-presenting cells (APCs) unlabeled HLA-A2/CD80-engineered K562 target-cells were used; K562 cells were pulsed with CMV pp65 peptide for activating conditions. K562 cell suspensions (5 Mio./ml) were kept on ice before pipetting into mounted sample chambers containing imaging buffer, and several minutes prior to adding T-cells to allow for settling. Cells were pipetted at a 1:2 ratio of target to T-cells. Imaging used the Sinc3 30×1000 (μm × nm) lattice-light-sheet and was set up as a single-track, dual-camera acquisition, directing DARPin-AF647 emission to camera 1 (red channel) using the SBS LP640 beam splitter and BP 570-620+LP655 emission filter; emission from K562 cells (green channel) used the BP 495-550/BP 570-620 filter before reaching camera 2. DARPin-AF647 (TCR) was excited at 640 nm (5-6% instrument power) with 10 ms/plane exposure; autofluorescence of APCs (K562) was excited at 488 nm (15%) with 10 ms/plane exposure. Volumetric sample scans were acquired with a 0.2 μm interval between slices over a 300 μm x-range (1051 slices/frame), yielding raw frames of 2048 × 228 pixels (optimal chip mode). The effective field of view was ∼300 μm × 33 μm (x × y) with a native voxel size of 0.145 μm × 0.145 μm × 0.2 μm (x, y, z). Sample scans were combined with time series, acquiring one sample scan every 60 seconds for total durations up to 10 minutes.

Raw data were processed in ZEISS ZEN (“Lattice Lightsheet” module). Volumes were deskewed using the “coverglass transformation” (linear interpolation). Processed data were resampled to isotropic 0.145 μm voxels (resulting in processed volumes of ∼329 μm × 300 μm × 16.4 μm) and examined as transformed 2D z-stacks and 3D maximum-intensity renderings (Precise mode). Cells of interest were cut out using the ROI tool, and sub-stacks of processed slices were created. 3D render series of frames of interest were created using the “Series” and “Positions List” option and exported as PNG images using the “Image Export” module. Final figures from exported images were created using FIJI/ImageJ (v1.54p).

## Acknowledgements

This work was supported by TU Wien and the European Union’s Horizon 2020 Marie Skłodowska-Curie COFUND doctoral program ENROL-Engineering for Life Sciences (grant 101034277), the Austrian Science Fund (FWF projects P 36923-B (E.S.), I 6611B (B.K. and M.B.)), and the WWTF (Project 2504476 (E.S.)).

