## Supplementary information for "A Minimally Perturbative DARPin Probe for Quantitative Fluorescence Imaging of the Human TCR-CD3 Complex"

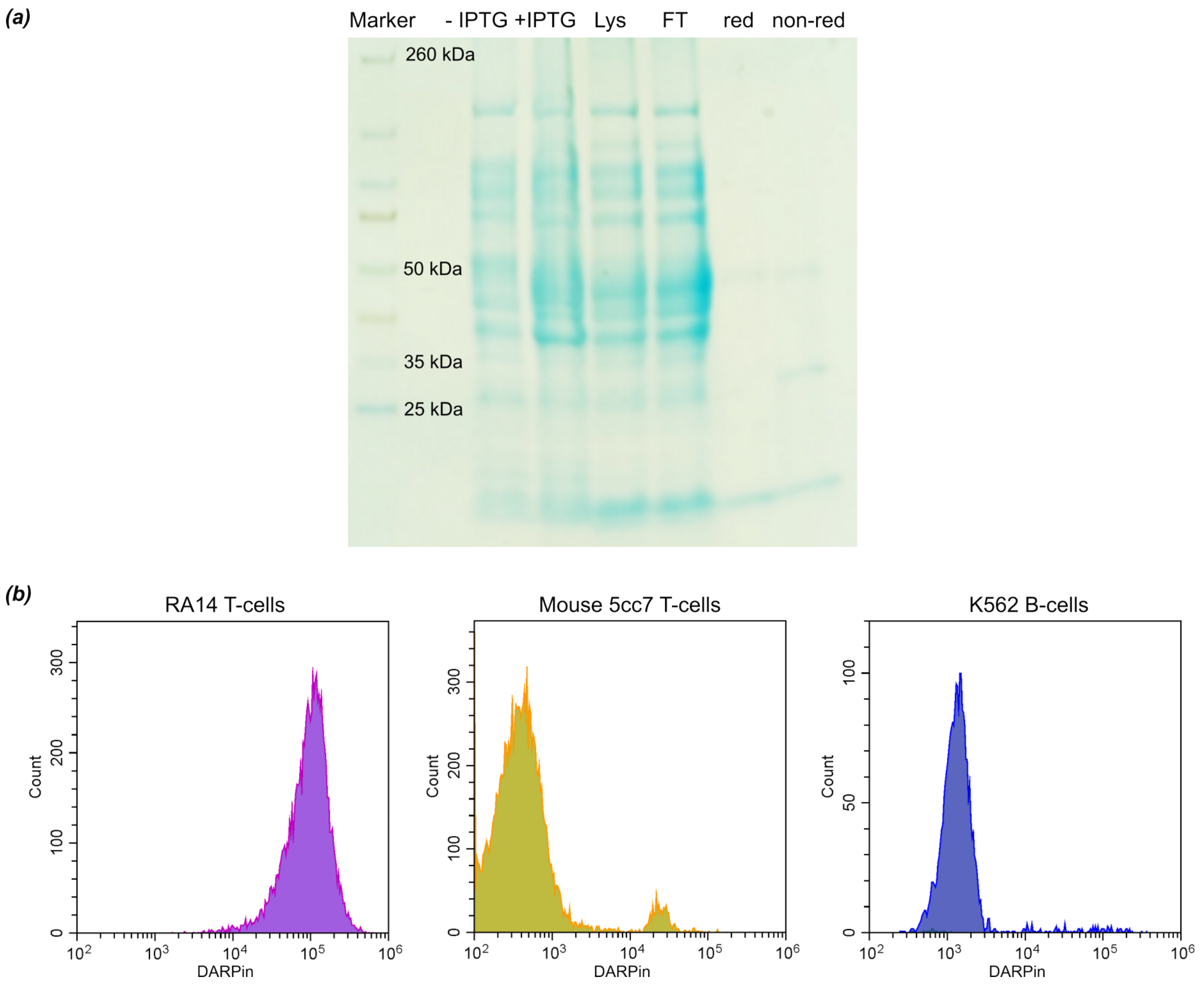


**Supplementary Figure 1. DARPin purification and binding specificity**

(*a*) Purification of the DARPin and SDS-PAGE analysis. Lanes: Marker of molecular weights, –IPTG (cell pellet prior to induction of protein expression), +IPTG (cell pellet at harvest following induction), Lys (bacterial lysate after sonication), FT (flow-through), red (final protein extract in reducing sample buffer, only monomer band is present); non-red (final protein extract in non-reducing sample buffer, covalent dimer and monomer bands are both visible). *(b)* DARPin binds negligibly to cells that do not express the human CD3 complex assessed with flow cytometry. Histograms illustrating binding of DARPin-AF647 to RA14 human T-cells (magenta, positive control), mouse 5cc7 T-cells (yellow), and human K562 B-cells (blue).

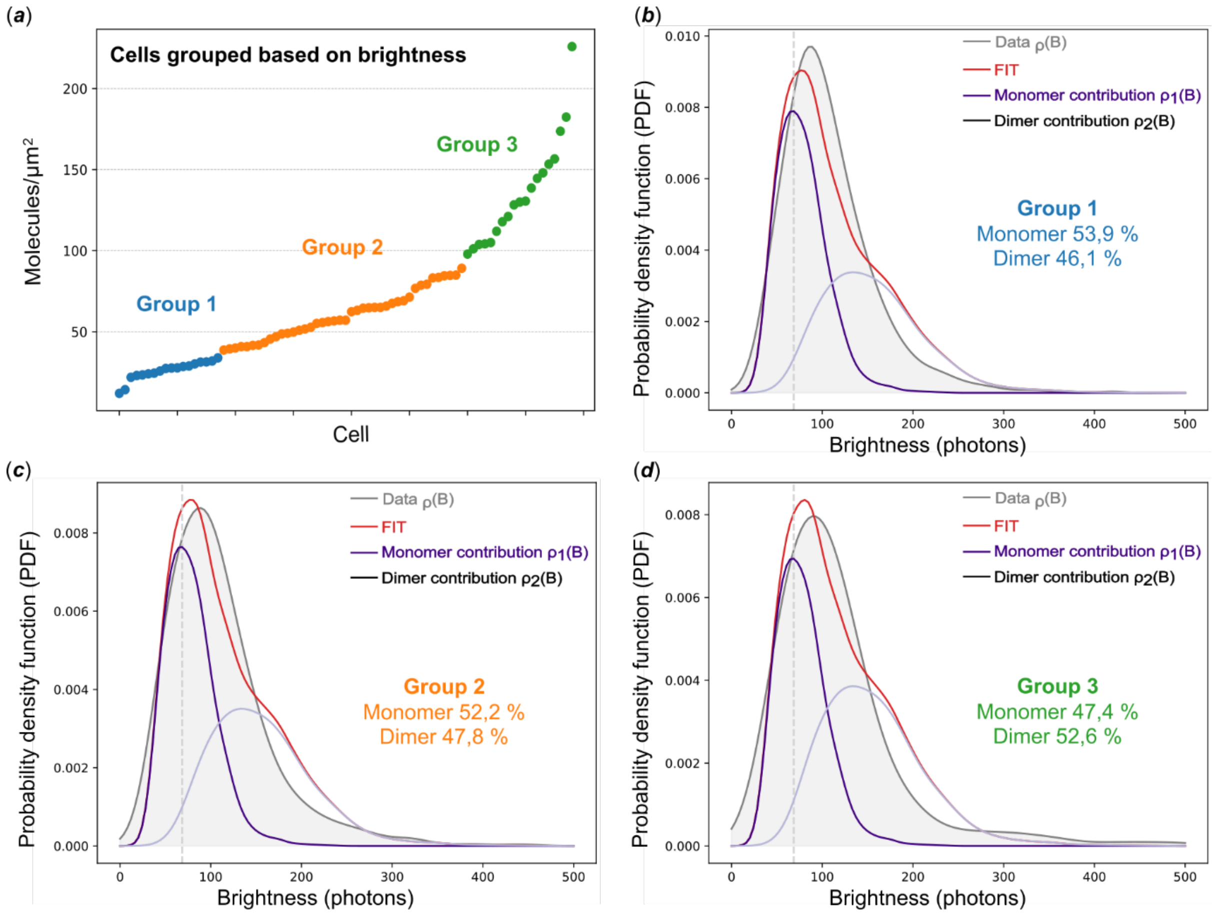


**Supplementary Figure 2. TOCCSL analysis stratified by DARPin labelling density.** (*a*) T cells were divided into three groups based on DARPin-AF647 surface density. TOCCSL brightness distributions were analyzed separately for each group, yielding monomer and dimer fractions. Results for group 1 (*b*), group 2 (*c*), and group 3 (*d*) show that the dimer fraction is independent of surface density, confirming that lower DARPin staining reflects lower TCR–CD3 expression rather than probe dissociation.

**Supplementary movie 1. DARPin-Cy3B_ICAM only.** Representative TIRF image of a DARPin-Cy3B-labeled RA14 T-cell on an SLB featuring ICAM only, recorded 5 min after initial contact. Scale bar, 10 µm.

**Supplementary movie 2. DARPin-Cy3B_pMHC-Alexa647_microclsuters.** Representative TIRF image of a DARPin-Cy3B-labeled RA14 T-cell interfaced with an SLB featuring pMHC-Alexa647 at 10 molecules/$\mu m^2$. Scale bar, 10 µm.

**Supplementary movie 3. DARPin-Cy3B_pMHC-Alexa647_cSMAC.** Representative TIRF image of a DARPin-Cy3B-labeled RA14 T-cell interfaced with an SLB featuring pMHC-Alexa647 at 10 molecules/$\mu m^2$ recorded 8min 30s after initial contact. Scale bar, 10 µm.
